# Therapeutic activation of endothelial sphingosine 1-phosphate receptor-1 by chaperone-bound S1P suppresses proliferative retinal vasculopathy

**DOI:** 10.1101/2022.06.14.496172

**Authors:** Colin Niaudet, Bongnam Jung, Andrew Kuo, Steven Swendeman, Edward Bull, Takahiro Seno, Reed Crocker, Zhongjie Fu, Lois E. H. Smith, Timothy Hla

**Affiliations:** Vascular Biology Program, Boston Children’s Hospital, Department of Surgery, Harvard Medical School, Boston, MA, USA; Department of Ophthalmology, Boston Children’s Hospital, Harvard Medical School, Boston, MA, USA

## Abstract

Sphingosine-1-phosphate (S1P), a bioactive lipid mediator that signals via G protein-coupled S1P receptors (S1PR), is required for normal vascular development. The role of this signaling axis in vascular retinopathies is largely unexplored. Here we show in a mouse model of oxygen-induced retinopathy (OIR) that endothelial overexpression of *S1pr1* suppresses while endothelial knockout (KO) of *S1pr1* worsens neovascular tuft formation. Furthermore, neovascular tufts are increased in *Apom* KO mice which lack HDL-bound S1P while they are suppressed in *Apom* TG mice which has more circulating HDL-S1P. These results suggest that circulating HDL-S1P activation of endothelial S1PR1 specifically suppresses proliferative retinal vasculopathy. Moreover, systemic administration of ApoM-Fc-bound S1P or a small molecule Gi-biased S1PR1 agonist suppressed neovascular tuft formation. Circulating HDL-S1P activation of endothelial S1PR1 may be a key protective mechanism to guard against neovascular retinopathies that occur not only in premature infants but also in diabetes and aging.

## Introduction

The retina, an extension of the central nervous system dedicated to light detection, is supplied by the specialized retinal and choroidal blood vessels. Pathological changes in these vascular beds represent a major cause of vision loss, and can occur either in the adult (i.e. diabetic retinopathy and age-related macular degeneration) or in infants suffering from retinopathy of prematurity (ROP). A common theme to such diseases is pathological angiogenesis (neovascularization) that follows tissue ischemia (Kermorvant-Duchemin et al., 2010). Inhibition of vascular endothelial growth factor (VEGF)-driven pathological angiogenesis has been the mainstay of therapeutic approaches for retinal neovascular diseases. However, this approach has limitations due to the requirement for intravitreal administration of VEGF neutralizing agents and the involvement of VEGF signaling in the maintenance of the choroidal vasculature and retinal neurons. Such constraints have prompted the search for alternative therapeutic approaches, particularly a treatment that inhibits neovascularization without affecting normal vascular development (Saint-Geniez et al., 2009; Kurihara et al., 2012; Bakri et al., 2014; Usui-Ouchi and Friedlander, 2019).

Sphingosine-1-phosphate (S1P), a bioactive sphingolipid, is required for vascular development (Mizugishi et al., 2005). In the developing vessels, S1P acts through its endothelial receptor S1PR1 to restrict VEGF-induced angiogenesis and induce vascular stability in multiple vascular beds (Chae et al., 2004; Gaengel et al., 2012; Jung et al., 2012; Ben Shoham et al., 2012). We recently showed that S1PR1-induced adherens junction formation suppresses VEGF-induced JunB induction in the primary vascular network, which allows Wnt-dependent organotypic specialization events to take place, thus maturing the retinal vasculature (Yanagida et al., 2020). The role of S1P signaling in the mature retinal vasculature is not known. Physiological S1P signaling is also affected by its carriers (named S1P chaperones) in blood, which assist in the activation of G-protein coupled receptors for S1P on target cells (Christoffersen et al., 2011). How S1P signaling impacts the endothelium in proliferative retinopathies is poorly understood. In oxygen-induced retinopathy (OIR), a mouse model that mimic the human ROP and other proliferative retinopathies (Smith et al., 1994), *Sphk2* constitutive knockout mice (Eresch et al., 2018), mice treated with a blocking antibody against S1P (Xie et al., 2009) as well as *S1pr2* KO mice (Skoura et al., 2007) were all partially protected from pathological neovascularization. These results suggest that S1P signaling via S1PR2, possibly in myeloid cells and pericytes, exacerbates the pathological phenotypes such as vascular leak, inflammation and endothelial proliferation in ROP. However, in humans, circulating S1P is decreased in severe ROP (Nilsson et al., 2021). To clarify the role of S1P in OIR, we studied mutant mice that lack or overexpress endothelial S1PR1 as well as circulating Apolipoprotein M (ApoM), which is the HDL-bound physiological S1P chaperone and studied retinal angiogenesis in the mouse OIR model. Our results show that circulating HDL-bound S1P and endothelial S1PR1 suppresses pathological neovascular tuft formation by inhibiting abnormal vascular leakage while enabling pericyte ensheathment. Systemic administration of recombinant ApoM-Fc-bound S1P or a small molecule G_i_-biased agonist of S1PR1 reduced neovascular tuft formation in OIR, suggesting that this pathway is a potential therapeutic target in neovascular retinal pathologies.

## Results and Discussion

### Endothelial S1PR1 suppresses neovascular tuft formation in OIR

S1PR1, the most abundant S1P receptor, is expressed in endothelial cells (EC) of the retina (Jung et al., 2012). To study the effect of S1PR1 signaling on pathological neovascularization that occurs in the OIR model, *S1pr1* overexpression in EC (*S1pr1* ECTG) was induced by tamoxifen from postnatal day 12 to 14 (P12-14). This regime was chosen because S1PR1 overexpression from P12-14 did not influence the normal development and maturation of primary or deep vascular plexuses (Fig. S1 A to E). This regime was then combined with exposure to hyperoxic stress from P7-12, which induces retinal vascular dropout and subsequent retinal hypoxia when mice are returned to a normoxic environment (relative hypoxia) from p12-17, with maximum neovascularization at P17 in WT mice, with resolution from P17-P21.

Analysis of droplet-based single-cell RNA sequencing (scRNA-seq) data from P17 retina (Binet et al., 2020) revealed that *S1pr1* is expressed at the highest level in the endothelium, while lower levels were also detected in the Müller glial cells. In addition, *S1pr1* is also the most abundant and frequently expressed among the three S1P receptors (*S1pr1, 2* and *3*) in retinal endothelium. OIR challenge did not alter S1PR1 expression at P14 or P17 (Fig. S1 F). *S1pr3* was detected in retinal pericytes whereas *S1pr2* expression was barely detectable in the retinal cells by scRNA-seq. The presence of S1PR1 protein in capillaries and neovascular tufts was confirmed by immunofluorescence staining of retinal sections post-OIR. As expected, *S1pr1* ECTG showed the strongest endothelial S1PR1 expression, whereas *S1pr1* ECKO mice were essentially devoid of S1PR1 immunoreactivity in EC (Fig. S1 G).

In the OIR model, *S1pr1* overexpression from P12-14 resulted in a strong decrease in the total area of neovascular tufts in P17 retinas (Fig. 1A and B). Interestingly, *S1pr1* ECTG mice also displayed significantly smaller individual neovascular tufts (Fig. 1B), which could also be observed on cross-sections (Fig. 1D). However, avascular areas were not altered by EC S1PR1 overexpression, indicating that normal revascularization was unaffected. The reduction in neovascular tufts in *S1pr1* ECTG retinas was observed as early as P15 (Fig. S1 H), suggesting that S1PR1 suppresses the formation of neovascular lesions.

**Figure 1.**
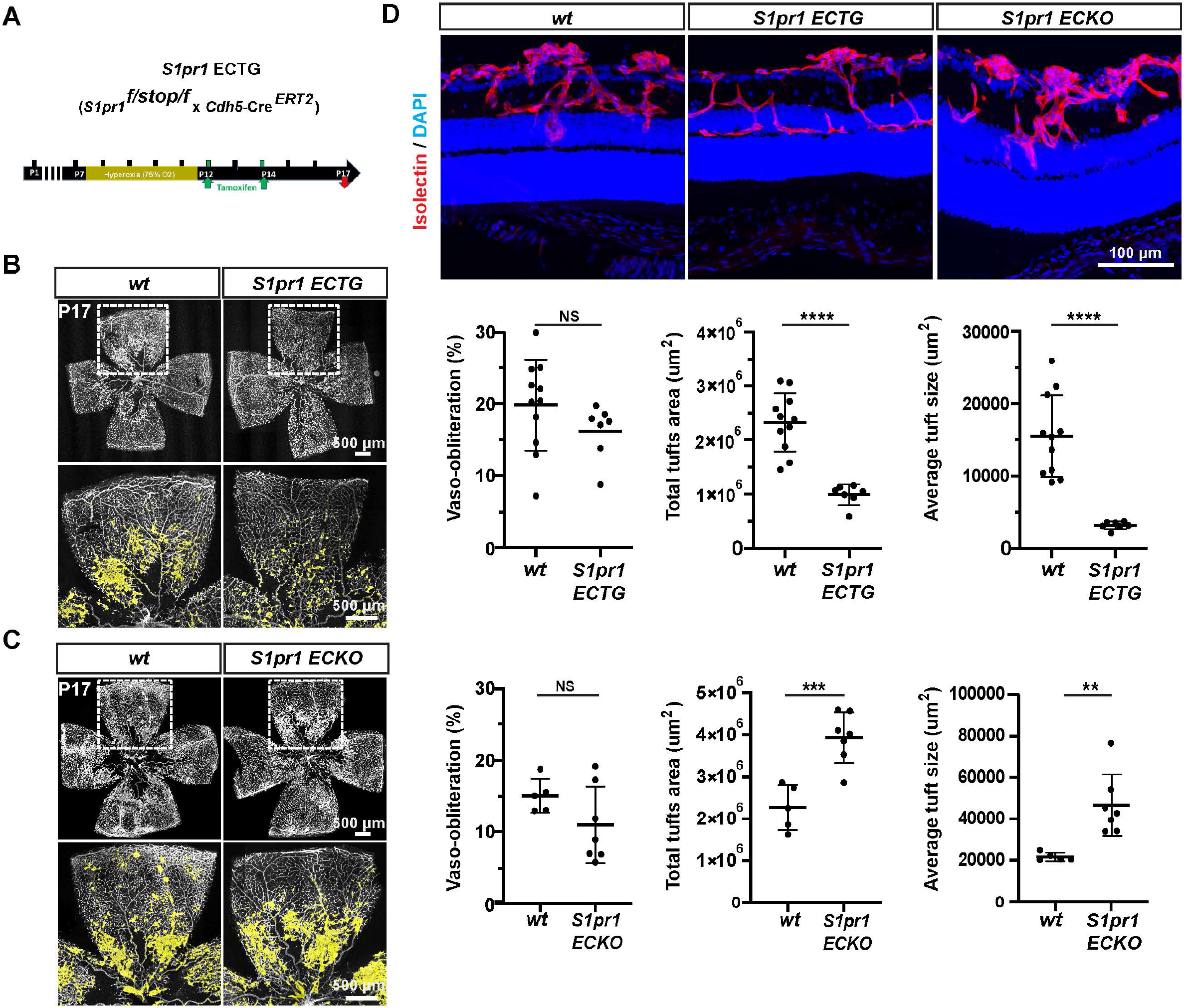
Endothelial S1PR1 signaling suppresses pathological neovascular tufts in OIR. (A) Strategy to induce S1PR1 expression in the endothelium post-OIR. *S1pr1* ECTG pups were exposed to 75% oxygen from P7 to P12. Upon return to normoxia, S1PR1 expression was induced by tamoxifen-injection at P12 and P14, and retinas were analyzed at P17 (B) Flat-mounted retinas from post-OIR WT and *S1pr1* ECTG pups at P17. Blood vessels are stained with isolectin (left) and pathological neovascular tuft area are highlighted in yellow. Avascular area, total and average neovascular tuft areas are quantified (right) (C) Flat-mounted retinas from post-OIR WT and *S1pr1* ECKO pups, at P17. Blood vessels are stained with isolectin and neovascular tufts are highlighted in yellow. Avascular area, total and average neovascular tuft areas are quantified. (D) Cross sections from post-OIR WT, *S1pr1* ECTG and *S1pr1* ECKO P17 pups. Blood vessels are stained with isolectin (red) and nuclei with Hoechst (blue). Data in B and C were analyzed by one-tailed Student’s *t* test. *, P < 0.05; **, P < 0.01; ***, P < 0.001; ****, P < 0.0001. A minimum of 5 pups were analyzed per group.

In contrast, genetic inactivation of *S1pr1* in the endothelium (*S1pr1* ECKO) at P12 resulted in an increase in the total area of neovascular tufts as well as an increased size of individual tufts at P17 in the OIR model (Fig. 1C and D). In both *S1pr1* ECTG and ECKO, the avascular area was not significantly changed, indicating that S1PR1 signaling in the endothelium does not influence the normal rate of re-vascularization. While this transgene induction regime did not impact vascular morphogenesis in normoxic retinas and the extent of the initial vaso-obliteration from P7-P12, it reduced the formation of neovascular lesions at P15 and P17. Overall, our results using gain- and loss-of function genetic models suggest that active S1PR1 signaling in the EC restrains the development of pathological angiogenesis in OIR, without affecting normal vascularization. These data also highlight the difference between S1PR1 agonism *vs*. anti-VEGF approaches, since the latter strategy blocked normal as well as pathological neovascularization (Tokunaga et al., 2014).

### Endothelial intrinsic function of S1PR1 affects vascular leakage, pericyte coverage and neovascular tuft resolution

OIR-induced vascular leakage was attenuated in *S1pr1* ECTG animals, as evidenced by fibrinogen staining that remained largely confined to the endothelial lumen in *S1pr1* ECTG tufts. By contrast, fibrinogen reached the basement membranes of WT tufts, and appeared more diffused in the retina of *S1pr1* ECKO (Fig. 2A). These results suggest that S1PR1 signaling counteracts the increased vascular permeability associated with OIR-driven hypoxia and other pathological processes. We also assessed the pericyte coverage of tufts. In WT and *S1pr1* ECTG, NG2 positive pericytes covered most of the surface of the neovascular tufts. In contrast, the pericyte coverage was much sparser in *S1pr1* ECKO tufts. Pericytes frequently lacked secondary processes and appeared detached from EC of neovascular tufts (Fig. 2B). It is likely that S1PR1 signaling in the EC regulates pericyte attachment in the retinal vasculature during OIR, as it was shown in coculture experiments with EC and mural cells (Paik et al., 2004).

**Figure 2.**
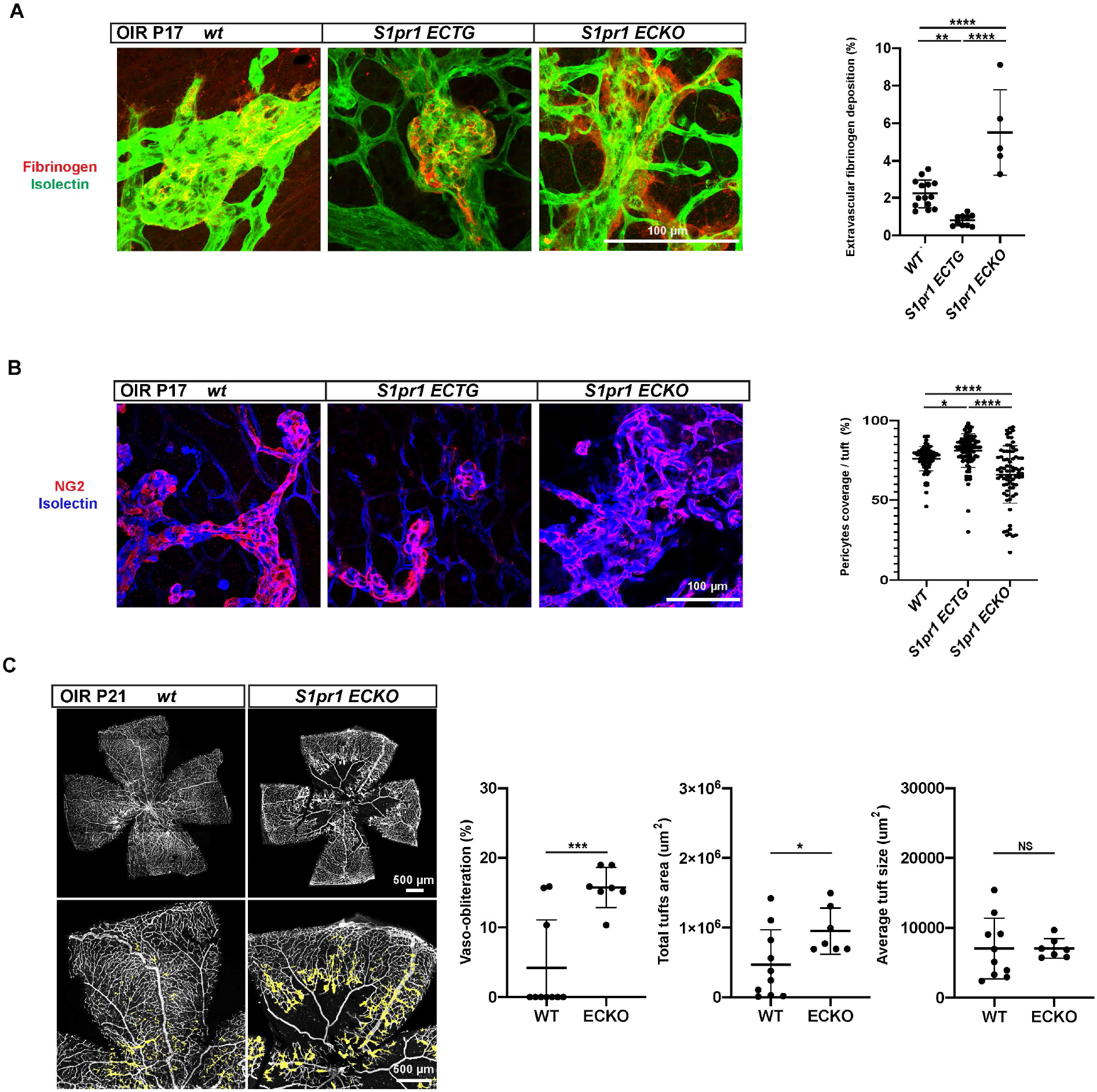
Endothelial S1PR1 attenuates vascular leakage and supports pericyte/EC interactions post-OIR. (A) Retinal flat mounts from post-OIR WT or *S1pr1* ECTG or ECKO P17 pups. Sites of vascular leakage were assessed by staining for fibrinogen (red), blood vessels are delineated by isolectin (green). Extravascular fibrinogen was quantified as described. (B) Pericytes stained by NG2 on the surface of isolectin-positive neovascular tufts at P17 post-OIR (left). Pericyte associated EC in neovascular tufts was quantified (right). (C) Flat-mounted retinas from post-OIR WT and S1pr1 ECKO pups at P21 stained with isolectin. Avascular area, total and average neovascular tuft areas are quantified. Data were analyzed by one-tailed Student’s *t* test. *, P < 0.05; **, P < 0.01; ***, P < 0.001; ****, P < 0.0001. At least 70 tufts coming from 3 different animals were assessed.

Recent studies suggested that neutrophils are involved in resolution of neovascularization in the OIR model (Binet et al., 2020). However, the number of retinal leukocytes, macrophages and neutrophils was similar in *S1pr1* ECTG retinas *vs*. WT counterparts (Fig. S2 A and B). The rate of neovascular tuft resolution was comparable regardless of S1PR1 expression level, when comparing *S1pr1* ECTG, ECKO and WT (Fig. 2C and Fig. S2 C) at days P19-21. *S1pr1* ECKO tufts, which are greatest at P17, were still evident at P21 whereas WT tufts have mostly resolved. In addition, neovascularization was greater in WT than in *S1pr1* ECTG counterparts at P19. Together, these results indicate that EC S1PR1 signaling regulated neovascular tuft formation and neovascular maintenance via EC intrinsic mechanisms. This signaling axis did not seem to impact leukocyte-dependent neovascular tuft resolution processes.

Taken together, these results indicate that S1PR1 signaling in the EC during OIR regulates pericyte/ EC interactions and vascular barrier function thereby impacting the extent of neovascularization. Since vascular leakage is critical for the inflammatory process, these results demonstrate that EC protective and anti-inflammatory functions of S1PR1 (Galvani et al., 2015; Christensen et al., 2016) are involved in the suppression of neovascular tuft formation in OIR.

### Therapeutic activation of S1PR1 suppresses neovascular retinopathy

To address whether S1P ligand bioavailability would impact neovascularization in OIR, we exposed mice in which the HDL-bound S1P is absent-i.e. *Apom* knockout (KO) mice to OIR (Christoffersen et al., 2011). At P17 when neovascularization is usually at its maximum, retinas from *Apom* KO exhibited severe pathological neovascularization, as compared to littermate controls (Fig. 3A). Conversely, mice constitutively overexpressing ApoM (*Apom* TG), which contain 7-9x more plasma ApoM than WT mice, had less retinal neovascular tuft formation at P17 after OIR (Fig. 3B). These data suggest that circulating HDL-bound S1P action on EC S1PR1 suppresses neovascular lesion formation in the OIR model, and established a proof of concept for ligand-based strategies.

**Figure 3.**
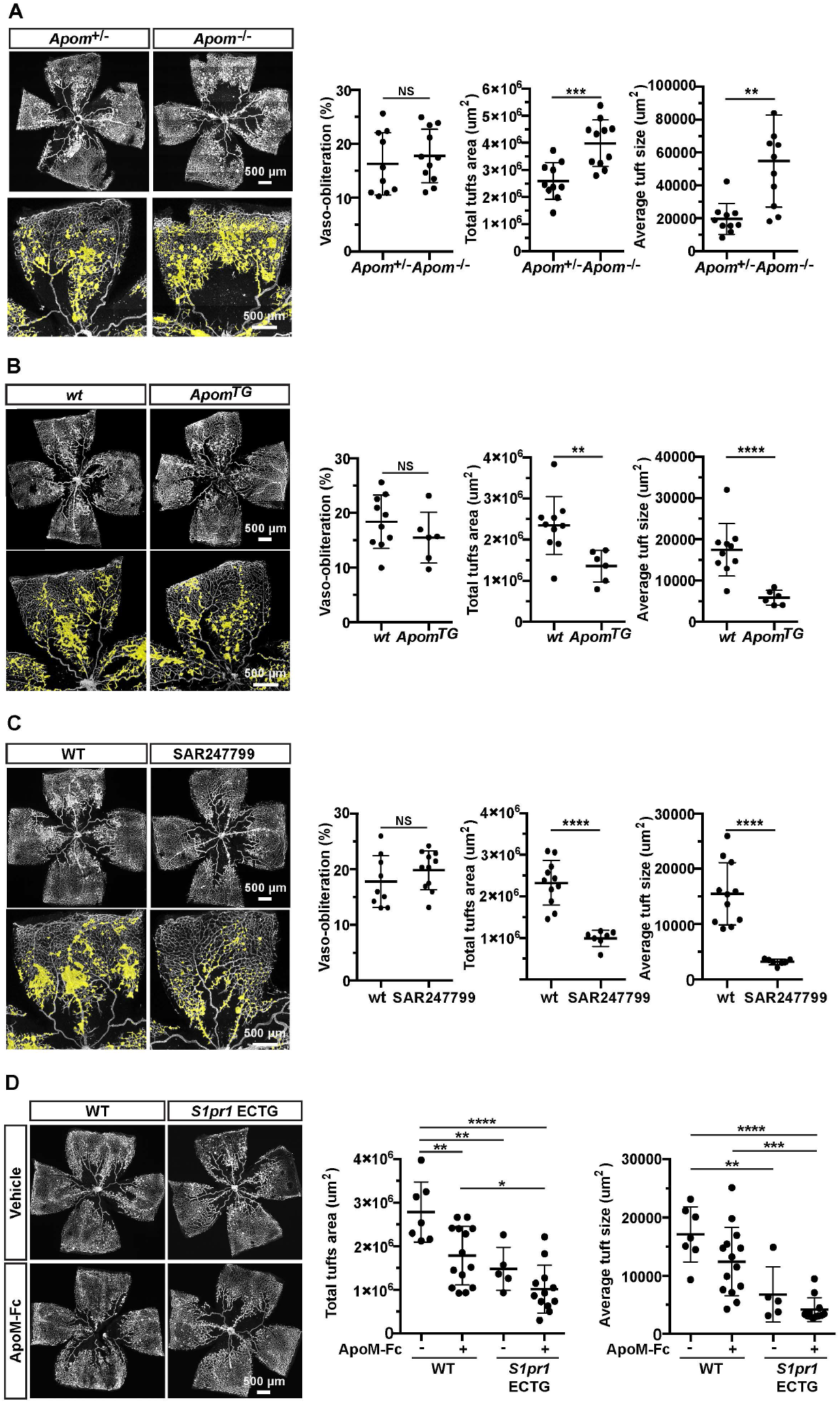
Systemic administration of S1PR1 agonists as a therapeutic strategy in retinopathy. (A) Flat-mounted retinas from post-OIR WT and *ApoM* KO pups at P17, stained with isolectin. Avascular area, total and average neovascular tuft areas are quantified. (n= 10 pups /group) (B) Flat-mounted retinas from post-OIR WT and *ApoM* TG pups at P17, stained with isolectin. Avascular area, total and average neovascular tuft areas are quantified. (n= 7 pups /group) (C) Flat-mounted retinas from SAR247799-treated, post-OIR WT pups at P17, stained with isolectin. Avascular area, total and average neovascular tuft areas are quantified. (n= 10 pups /group) (D) Flat-mounted retinas from ApoM-Fc-treated, post-OIR WT or S1pR1 ECTG pups at P17, stained with isolectin. Avascular area, total and average neovascular tuft areas are quantified. (n= 6 pups /group) Data in B and D were analyzed by one-tailed Student’s *t* test. *, P < 0.05; **, P < 0.01; ***, P < 0.001.

HDL-S1P selectively activates the EC S1PR1 and does not influence lymphocyte trafficking from secondary lymphoid organs and thymus. This is thought to be achieved via biased signaling of S1P on EC S1PR1 (Galvani et al., 2015; Poirier et al., 2020). For example, HDL-S1P induces sustained EC barrier function compared to albumin-S1P or small molecule S1PR1 modulators which induce robust ß-arrestin-dependent S1PR1 internalization (Wilkerson et al., 2012; Christensen et al., 2016). In contrast, the large size of HDL-S1P likely prevents it from modulating lymphocyte S1PR1 in secondary lymphoid organs or thymus, which is needed for modulation of lymphocyte trafficking.

Recently, a small molecule G_i_-biased agonist of S1PR1 was shown to activate EC S1PR1 and suppress inflammatory processes without inducing lymphopenia. In humans, this compound, SAR247799, enhanced NO-dependent cardiac reperfusion in diabetic subjects in a human phase 1 clinical study (Bergougnan et al., 2021) without inducing substantial lymphopenia. We tested the ability of SAR247799 to modulate retinal vascular tuft formation in ROP. SAR249799 was injected for five consecutive days to pups post oxygen-exposure in OIR. In OIR, oxygen exposure (P7-P12) causes loss of vessels and ischemia, thus initiating the neovascular phase (P12-P17). We found that neovascular tuft formation was suppressed by systemic SAR247799 administration (Fig. 3C). This suggests that G_i-_biased agonism of S1PR1 leads to the suppression of neovascular tuft formation in OIR.

Since the high circulating HDL-S1P levels phenocopied EC *S1pr1* ECTG in OIR, we explored the hypothesis that systemic administration of ApoM-bound S1P may be a viable therapeutic strategy to activate endothelial S1PR1 during ROP and other proliferative retinopathies. An injectable recombinant ApoM-Fc fusion protein (Swendeman et al., 2017) was administered post oxygen-exposure in OIR (at initiation of neovascularization) to WT and *S1pr1* ECTG mice. ApoM-Fc retains its binding capacity for S1P, and leads to sustained endothelial signaling, without affecting lymphocyte egress (Swendeman et al., 2017). Systemic administration of recombinant ApoM-Fc at P13 and P15 post oxygen-exposure in OIR reduced neovascular tufts formation, compared to vehicle injected littermates at P17. Enhancing both ligand and receptor (in ECTG) led to an additive reduction in neovascular tuft formation (Fig. 3 D).

From a therapeutic perspective, our results show that systemic treatment with S1PR1 agonists protects the endothelium against OIR without suppressing normal re-vascularization. S1PR1 activation could therefore represent an important complement to intravitreal, VEGF-targeted strategies which suppress both neovascularization as well as normal re-vascularization. In this regard, the ability of S1PR1 to restore endothelial functions could be important in patients refractory to anti-VEGF treatments, and all patients with retinopathy at risk for suppression of normal vascular maintenance and development. On the other hand, much remains to be understood regarding the bioavailability of S1P during ROP. The mechanisms explaining the systemic changes in plasma S1P levels during ROP are ill-defined, while the role played by lipids in retinopathies, and more specifically ROP, is increasingly acknowledged (Fu et al., 2015). Moreover, intra-retinal variations in S1P synthesis and export also require further exploration, as with other lipids (Gantner et al., 2019). An alternative to S1P therapy could be to increase S1PR1 levels in retinal endothelium via gene therapy approaches (Cepko, 2012), although the effects might be less immediate than through ligand administration. Taken together, data from this study suggests that activation of the S1PR1 pathway may be an attractive candidate for treating proliferative retinopathies at large.

## Materials and methods

### Animals

All mouse experiments were approved by Institutional Animal Care and Use Committee of Boston Children’s Hospital. Transgenic mouse models used in this study are following: *S1pr1*^flox/flox^ (a kind gift of Dr. Richard Proia, NIDDK, NIH), *S1pr1* ^f/stop/f/ f/stop/f^ (Jung B, 2012), *Cdh5*-*Cre*^*ERT2*^ (a kind gift of Dr. Ralf Adams, Max Planck Institute, (Sörensen et al., 2009)), *Apolipoprotein M* transgenic (*Apom* TG) and *Apom* knockout (KO) mice (Christoffersen et al., 2011). For tamoxifen inducible, endothelial-specific induction or deletion of *S1pr1* gene in mice, either *S1pr1* ^f/stop/f/ f/stop/f^ or S1pr1 ^flox/flox^ mouse was crossed to *Cdh5-Cre*^*ERT2*^ that yielded *S1pr1* ^f/stop/f/ f/stop/f^; *Cdh5-Cre*^*ERT2*^ (herein referred to as *S1pr1* ECTG) and *S1pr1* ^flox/flox^; *Cdh5-Cre*^*ERT2*^ (herein referred to as *S1pr1* ECKO), respectively.

### Mouse experiments

Oxygen-induced retinopathy (OIR) was induced according to the protocol described previously (Smith et al., 1994). Briefly, pups at P7 and their nursing dam were transferred to a chamber (A-30274-P, Biospherix) with an oxygen concentration maintained at 75% (ProOx Model 110, Biospherix) till P12. When the pups were returned to room oxygen levels (21%), 150 ug tamoxifen (Sigma-Aldrich, CAT#T5648) dissolved in corn oil (Sigma-Aldrich CAT#C8267) via intraperitoneal injection was administered 2 times (at P12 and P14) in order to induce or delete *S1pr1* in mice. Both males and females were used, and littermates that do not bear *Cdh5-Cre*^*ERT2*^ gene were used as controls. Mice were sacrificed and eyes were collected at the time indicated.

To address pharmacological S1PR1 agonism upon OIR challenge, mice were given intraperitoneal injection of ApoM-Fc/S1P (100 µg/ mouse, 4 mg/Kg) at P13 and P15. SAR247799 (30 mg/Kg), S1PR1 specific agonist, was dissolved in saline, and administered via an intraperitoneal route for 5 consecutive days (from P12 to P16). Mice were sacrificed at P17 for retinal analysis. Vehicle controls were included within the littermates per experiment.

### Immunofluorescence staining

Whole mount staining of retinas was performed as previously described (Gaengel et al., 2012, Jung et al., 2012). Briefly, eyes were post-fixed in 4% PFA in PBS at room temperature for 30 min and retinas were dissected. For cryosections, eyes were cryoprotected in 30% sucrose in PBS overnight at 4°C, embedded in a 1:1 mixture of 30% sucrose PBS:OCT over dry ice and then sectioned at 30 µm intervals using a cryostat (Leica Biosystems). Retinas were then permeabilized in PBLec (1% Triton X-100 100 µM CaCl_2_, 100 µM MgCl_2_, MnCl_2_ in PBS (pH6.8)), at room temperature for 30 min, blocked with 1% Bovine Serum Albumin (BSA, Sigma-Aldrich, CAT#A6003)/ PBLec at room temperature for 30 min. Primary antibodies were diluted in 1% BSA/ PBLec and incubated at 4°C for overnight. After thorough washing in PBLec, fluorophore-conjugated or secondary antibodies were added to retinal tissue. Both primary and secondary antibodies used were anti-CD31 (1:300, R&D systems, CAT#AF3628), anti-collagen IV (Biorad), anti-ERG (1:300, Abcam, CAT#ab92513), anti-fibrinogen (1:500, Accurate Chemical, CAT#YNGMFBG), anti-CD45 (1:300, R&D systems, CAT#AF114-SP), anti-GFAP(1:300, Dako, CAT#ZO334), anti-NG2 (1:300, Millipore, CAT#MAB5320), anti-S1PR1 (H60, sc 25489, Santa Cruze biotechnology), Alexa Fluor 488-conjugated anti-ERG (1:300, Abcam, CAT#ab196374), Alexa Fluor 647-conjugated anti-ERG (1:300, Abcam, CAT#ab196149), Cy3-conjugated anti-aSMA (Sigma-Aldrich, CAT#C6198), Alexa Fluor 488- (1:500, CAT#I21411), Alexa Fluor 568- (1:500, CAT#I21412), or Alexa Fluor 647- (1:500, CAT#I32450) conjugated isolectin GS-IB4 (all from Thermo Fisher Scientific). Retinas were mounted using Fluoromount-G slide mounting medium (Southern Biotech, CAT#0100-01).

### Confocal Imaging

Images were acquired using a LSM810 confocal microscope (Zeiss) equipped with an EC Plan-Neofluar 10x/0.3, a Plan-Apochromat 20x/0.8, a Plan-Apochromat 40x/1.4 Oil DIC, or a Plan-Apochromat 63x/1.40 Oil DIC objective. Images were taken using Zen2.1 software (Zeiss), and processed and quantified with Fiji (NIH). Figures were assembled using Adobe Photoshop and Illustrator. Tufts, avascular and total retina areas were quantified with ImageJ software (http://rsb.info.nih.gov/ij/) (Connor et al., 2009). For leakage quantification, one retinal quadrant was imaged for a minimum of 5 animals per genotype, and the surface of fibrinogen-positive stain outside the isolectin-positive vascular area was quantified over the total retinal surface. For quantification of pericytes-coverage of neovascular tufts, images were taken at high resolution (40X) and over 70 tufts from three different animals per genotype were analyzed. The pericytes coverage was defined as the ratio of NG2-positive surface over the endothelial surface of neovascular tufts, delineated by isolectin-positive staining.

### FACS analysis

Retinas were dissected from freshly collected eyes and digested with LiberaseTM (Sigma-Aldrich, CAT#5401127001, 0.26 U/mL) and deoxyribonuclease I (Sigma-Aldrich, CAT#D4527, 10 mg/mL) in PBS (1 mL per one retina) at PBS at 37°C for 30 min. The reaction was stopped by adding SVF (final 1%) and centrifuged at 400 × g at 4°C for 10 min. The pellet was further washed once with FACS buffer (0.5% BSA/0.5 mM EDTA in PBS) and stained with APC-conjugated anti-mouse CD45 (1:300, BioLegend, CAT#103112), BV510-conjugated anti-mouse CD11b (1:300, Biolegend, #CAT101263) and PE-conjugated anti-mouse Ly6G (1:300, BD Pharmingen, #CAT5551461) antibodies for 1 hour on ice. The retinal cells were washed twice and treated with DAPI to exclude dead cells. CD31+/CD45-/TER119-cells were analyzed using BD FACSAria™ II (BD Biosciences).

### Single-cell RNAseq analysis

Data were accessed from NCBI’s Gene Expression Omnibus (accession no. GSE150703 and GSE141440). Downstream processing of the gene expression matrix was performed using the “Seurat” R package. Clustering followed by marker gene analysis enabled annotation of canonical retina cell types. Differences in gene expression frequency and intensity are visualized using the DotPlot and FeaturePlot functions.

### Statistical analysis

Data are expressed as mean ± SD. Statistical analyses were performed using GraphPad Prism software v.8.0. In datasets containing two distinct groups, statistical comparisons were performed with the Student’s t-test, and P<0.05 was considered statistically significant. In dataset containing three distinct groups, statistical comparisons among groups were performed using one-way ANOVA followed by Tukey’
ss post-hoc test and P<0.05 was considered statistically significant. On the figures, the error bars represent SD, and P<0.05 is represented as *, P<0.005 as **, P<0.0005 as ***, P<0.00005 as **** and “ns” stands for “no significant difference”. Number of animals represent biological replicates.

## Acknowledgements

The authors thank Dr Elizabeth Moran for initial experiments with ApoM-Fc. This work is supported by NIH grants RO1-EY031715 and R35-HL135821 to TH.

## Figure legends

**Figure S1.**
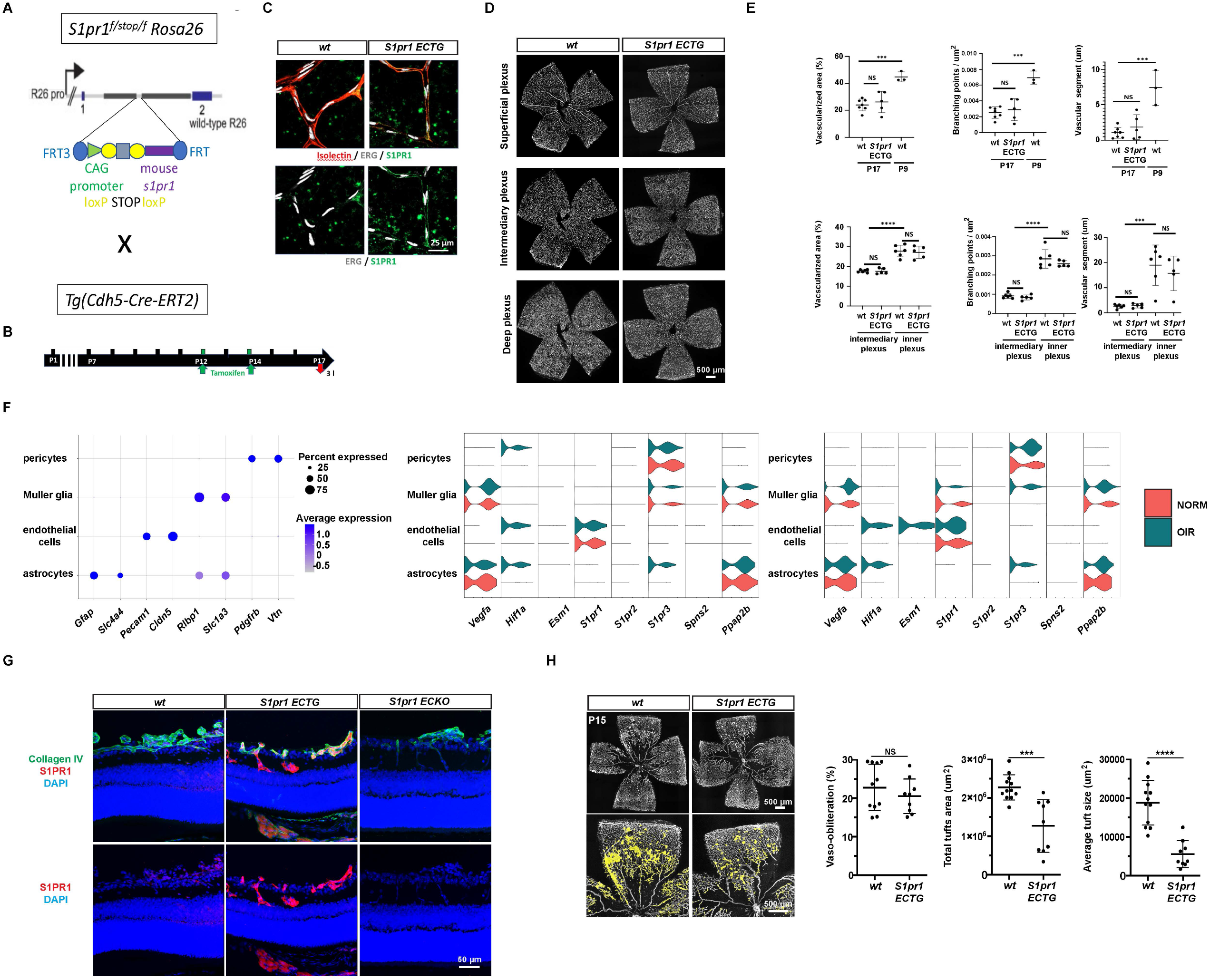
S1PR1 signaling in the retinal EC influences neovascular tuft development in OIR. (A) Schematic representation of the breeding strategy to establish *S1pr1* ECTG: females carrying the *(ROSA)26Sortm1(CAG-S1pr1)* transgene were crossed to *Cdh5-Cre*^*ERT2*^ Cre males. (B) Strategy to induce expression of S1PR1 in post-natal endothelium. *S1pr1* ECTG pups were given tamoxifen at P12 and P14, and retinas were analyzed at P17. (C) Flat-mounted retinas from post-OIR WT and *S1pr1* ECTG pups at P17. High magnification pictures of superficial capillaries showing S1PR1 induction in the *S1pr1* ECTG. (D) Flat-mounted retinas from post-OIR WT and *S1pr1* ECTG pups at P17. Retinal vasculature parameters were stained with isolectin and images of superficial (top panel), intermediary (middle panel) and deep (lower panel) plexuses are shown. (E) Quantification of morphometric parameters in between retinas from WT and *S1pr1* ECTG. Retinas from P9 WT mice (not shown) were used as controls. (F) Dot plot representing expression level and frequency of cell types markers among non-neuronal retinal cells at P14 and P17 in OIR (G) Volcano plot showing expression level and frequency of OIR-induced genes (left) and S1P-related genes (right) among non-neuronal retinal cells at P14 (left) and P17 (right) in OIR. *S1pr4* and *S1pr5* expression was limited to a low number of endothelial cells. (H) Cross sections from OIR WT, *S1pr1* ECTG and *S1pr1* ECKO P17 pups stained for S1PR1 (red) Blood vessels are delineated by collagen IV (green) and nuclei are stained with Hoechst (blue). (I) Flat-mounted retinas from post-OIR WT and *S1pr1* ECTG pups at P15. Blood vessels are stained with isolectin (left). Avascular area, total neovascular tuft area and average neovascular tuft size are quantified (right). Data in E were analyzed by one-tailed Student’s *t* test. *, P < 0.05; **, P < 0.01; ***, P < 0.001; ****, P < 0.0001. A minimum of 3 pups per group were analyzed.

**Figure S2.**
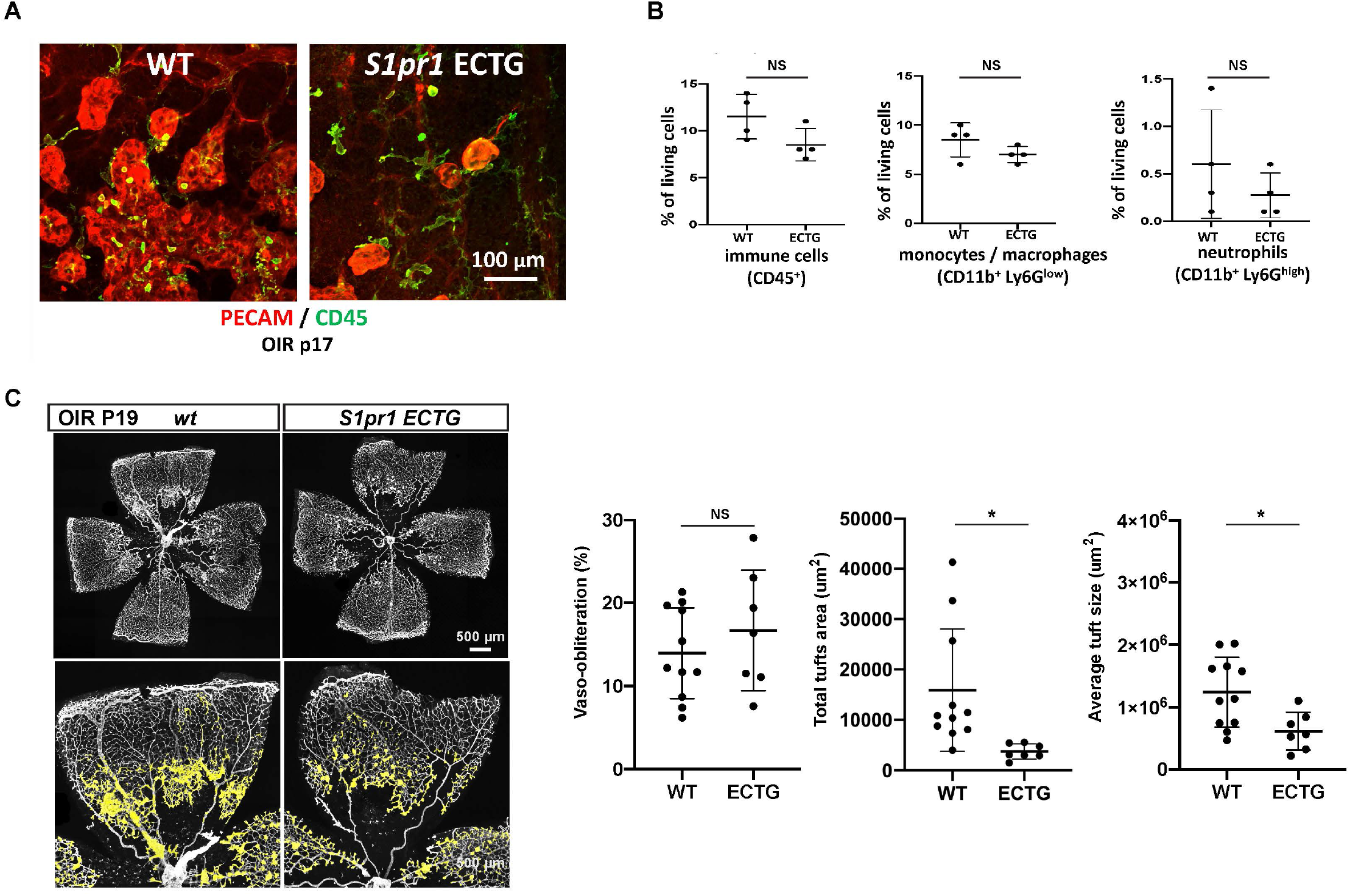
Leukocytes and neovascular tuft phenotypes during resolution phases of OIR. (A) Flat mount view of neovascular tufts from WT or *S1pr1* ECTG at P17, stained for immune cells (CD45), and blood vessels (isolectin). (B) FACS on retinal single-cell preparations. Quantification of immune cells by FACS in post-OIR retinas from WT and *S1pr1* ECTG at P17. Total immune cells (CD45 positive, left panel), monocytes/macrophages (CD11b positive, middle panel) and neutrophils (CD11b and Ly6G double positive, right panel) are presented. 3 pups or more were analyzed per group. (C) Flat-mounted retinas from post-OIR WT and *S1pr1* ECTG pups at P19 stained with isolectin. Avascular area, total and average neovascular tuft areas are quantified. Data were analyzed by one-tailed Student’s *t* test. *, P < 0.05. A minimum of 3 pups per group were analyzed.

